# Statistical Methods for Microrheology of Airway Mucus with Extreme Heterogeneity

**DOI:** 10.1101/2023.11.20.567244

**Authors:** Neall Caughman, Micah Papanikolas, Matthew Markovetz, Ronit Freeman, David B. Hill, M. Gregory Forest, Martin Lysy

## Abstract

The mucus lining of the human airway epithelium contains two gel-forming mucins, MUC5B and MUC5AC. During progression of cystic fibrosis (CF), mucus hyper-concentrates as its mucin ratio changes, coinciding with formation of insoluble, dense mucus flakes. We explore rheological heterogeneity of this pathology with reconstituted mucus matching three stages of CF progression and particle-tracking of 200 nm and 1 micron diameter beads. We introduce statistical data analysis methods specific to low signal-to-noise data within flakes. Each bead time series is decomposed into: (i) a fractional Brownian motion (fBm) *classifier of the pure time-series signal*; (ii) high-frequency static and dynamic noise; and (iii) low-frequency deterministic drift. Subsequent analysis focuses on the denoised fBm classifier ensemble from each mucus sample and bead diameter. Every ensemble fails a homogeneity test, compelling clustering methods to assess levels of heterogeneity. The first binary level detects beads within vs. outside flakes. A second binary level detects within-flake bead signals that can vs. cannot be disentangled from the experimental noise floor. We show all denoised ensembles, within- and outside-flakes, fail a homogeneity test, compelling additional clustering; next, all clusters with sufficient data fail a homogeneity test. These levels of heterogeneity are consistent with outcomes from a stochastic phase-separation process, and dictate applying the generalized Stokes-Einstein relation to each bead per cluster per sample, then frequency-domain averaging to assess rheological heterogeneity. Flakes exhibit a spectrum of gel-like and sol-like domains, outside-flake solutions a spectrum of sol-like domains, painting a rheological signature of the phase-separation process underlying flake-burdened mucus.

## Introduction

We first briefly recall the experimental and data-analysis protocols of Passive Particle-Tracking Microrheology (PPTM) [1, 2, 3]. Microbeads are embedded in a sample and their *position time series*,

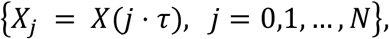

are extracted using particle-tracking microscopy, where *X*(*t*) is the continuous-time trajectory of a given microbead, *τ* is the *experimental lag time between recorded positions of the microscope*, and *Nτ* is the *total tracking time*. We enforce uniformity of *τ* and *N* for each tracked bead for statistical robustness of data inference. From the time series data, one then calculates increment statistics, specifically the time-averaged mean-squared-displacement of each tracked bead [23,24,25], *MSD*_*X*_, computed for all lag times *nτ*:

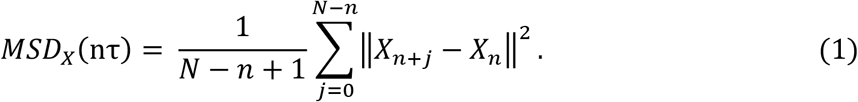

Positions *X*_*j*_ have dimension *d*, typically *d* = 2 with beads tracked in distinct focal planes.

*Pure medium-induced increments and MSD statistics, free from experimental noise and tracking error, are inserted into the Generalized Stokes-Einstein Relation (GSER)*, mapping from increments at lag times *nτ* to dynamic moduli at frequencies *ω* = 1*⁄nτ* The GSER can be applied to data of *individual tracked beads* of radius *r*, Equation (2), or to the *ensemble-averaged* MSD, Equation (3). The elastic, *G*^′^(*ω*), and viscous, *G*^′′^(*ω*), dynamic moduli of the medium surrounding individual beads (2) or a randomly sampled medium (3) are given by:

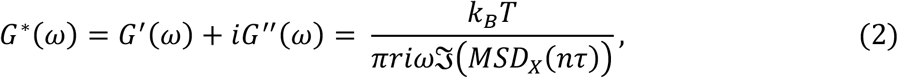

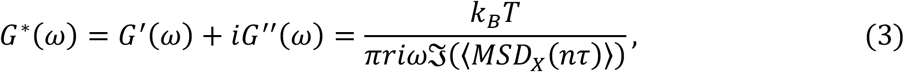

where ℑ(·) is the Fourier transform and *G*^*∗*^(*ω*) is the complex shear modulus. *Equation* (3) *is the standard formula employed for presumed homogeneous materials, where* ⟨· ⟩ *denotes the ensemble average over all tracked beads*. PPTM has been indispensable for soft complex fluids with extremely low yield thresholds and low volume availability, such as human bronchial epithelial (HBE) mucus (cf. [4-15, 21, 22]).

In addition to previously documented increases in mucus concentration [9,12], cystic fibrosis (CF) mucus networks reconfigure to contain [15,22] dense, raft-like, structures, referred to as *flakes*. In advanced stages of CF, up to 50% of the gel-forming mucins (MUC5B and MUC5AC) in airway mucus is sequestered within flakes that do not swell nor dissolve in airway surface salt-water solvent. Advanced CF mucus can therefore be represented, at a coarse scale, as a two-phase mixture: a dense phase of mucin-rich flakes, and a more dilute phase consisting of the remaining mucins. Additional scales of spatial heterogeneity potentially exist within the two coarse phases for any given sample [22]: the dilute phase may vary in mucin concentration; flake lateral dimensions range from 1-100 microns (μm), with unknown free-volume pore-size distributions that govern admissibility of passive probes added to samples. For this study, we disperse 200 nm and 1 μm diameter beads in separate sub-samples of the same reconstituted HBE mucus. As expected, only a few beads can be incorporated into dense flakes, and their diffusive mobility is hindered, sometimes sufficiently low to be indistinguishable from immobile beads tracked by the instrumentation, i.e., the tracked bead signal is sometimes entangled in the noise floor.

Heterogeneity and insufficient random sampling of flake-burdened mucus samples impose hard constraints on the form of the Generalized Stokes-Einstein Relation (GSER) one should use. The inherently low bead numbers within flakes further limit inference of rheology. Ensemble averaging of bead MSDs in the GSER is justified by an ergodicity assumption for homogeneous materials: ensemble averaging of bead MSDs in the time domain is equivalent to frequency-domain averaging of the Fourier-transformed MSDs, further assuming a random sampling of the material. These assumptions are strongly violated for flake-prevalent mucus [15,22].

To overcome these limitations, we proceed in the following steps:

- *Experimental*: Reconstitute bulk samples of HBE mucus to match the MUC5B:MUC5AC ratio during three progressive stages of CF [34]. Disperse 200 nm and 1 μm diameter beads within sub-samples from the same bulk samples. Use microscopy to record the position times series of beads per diameter and per sub-sample, for all 3 samples.
- *Filtering of the data*. Apply a proximity filter to ensure beads are isolated and not within 5 diameters of another bead to avoid bead-bead interactions and cross-correlations, cf. [17,17,18]. Filter any time series with missing increments.
- Given that within-flake beads often have mobilities on the order of the noise floor, *use the sophisticated denoising procedure* of [16], generalized in [33] and here *to analyze low mobility data from beads in flakes*. Explore several classifier metrics of individual bead time series and seek the optimal metric for the experimental data.
- *Use synthetic simulated data* to replicate features of the experimental data, in particular, heterogeneous tracked particle time series, extremely low mobility and signal-to-noise (SNR) time series, and then *to identify an optimal classifier metric* (described in detail below).
- *With the optimal classifier metric, hierarchically cluster the experimental datasets*. 1^st^, distinguish beads outside and within flakes. (The accuracy of this automated task is experimentally tested by visual assessment of beads outside and inside flakes, with 209/212 (98.58%) beads correctly identified.) 2^nd^, distinguish whether tracked beads within flakes can be confidently disentangled from the noise floor. 3^rd^, test whether the ensembles of denoised within-flake and outside-flake beads, per reconstituted sample, are statistically homogeneous. (N.B. All fail the homogeneity test.) 4^th^, given that within-flake and outside-flake ensembles are non-homogeneous, apply a clustering algorithm to the within-flake and outside-flake datasets for the three bulk samples for each bead diameter. 5^th^, test homogeneity of all individual clusters containing at least 4 beads. (N.B. All clusters fail the homogeneity test.)
- Given non-homogeneity of each cluster, *apply the GSER to single beads in each cluster and then cluster average in frequency space to infer the dynamic moduli of all denoised data*
- *clusters within and outside of flakes, for 200 nm and 1* μ*m diameter beads. Assess the resulting rheological heterogeneity* between clusters within and outside of flakes, for each probe size.
- *Assess probe length-scale dependence in the dynamic moduli* from the 200 nm and 1 μm bead diameter results for all three mucin-ratio samples. Make *preliminary inferences of poresize distributions of flakes* for the three CF-like mucin-ratio samples and summarize *future steps to make inferences of pore-size distributions rigorous*.

Figure 1 shows the mucosal flakes with individual beads numbered and highlighted. Here we see the vast difference in flake size as well as the number of beads within different flakes.

**Figure 1:**
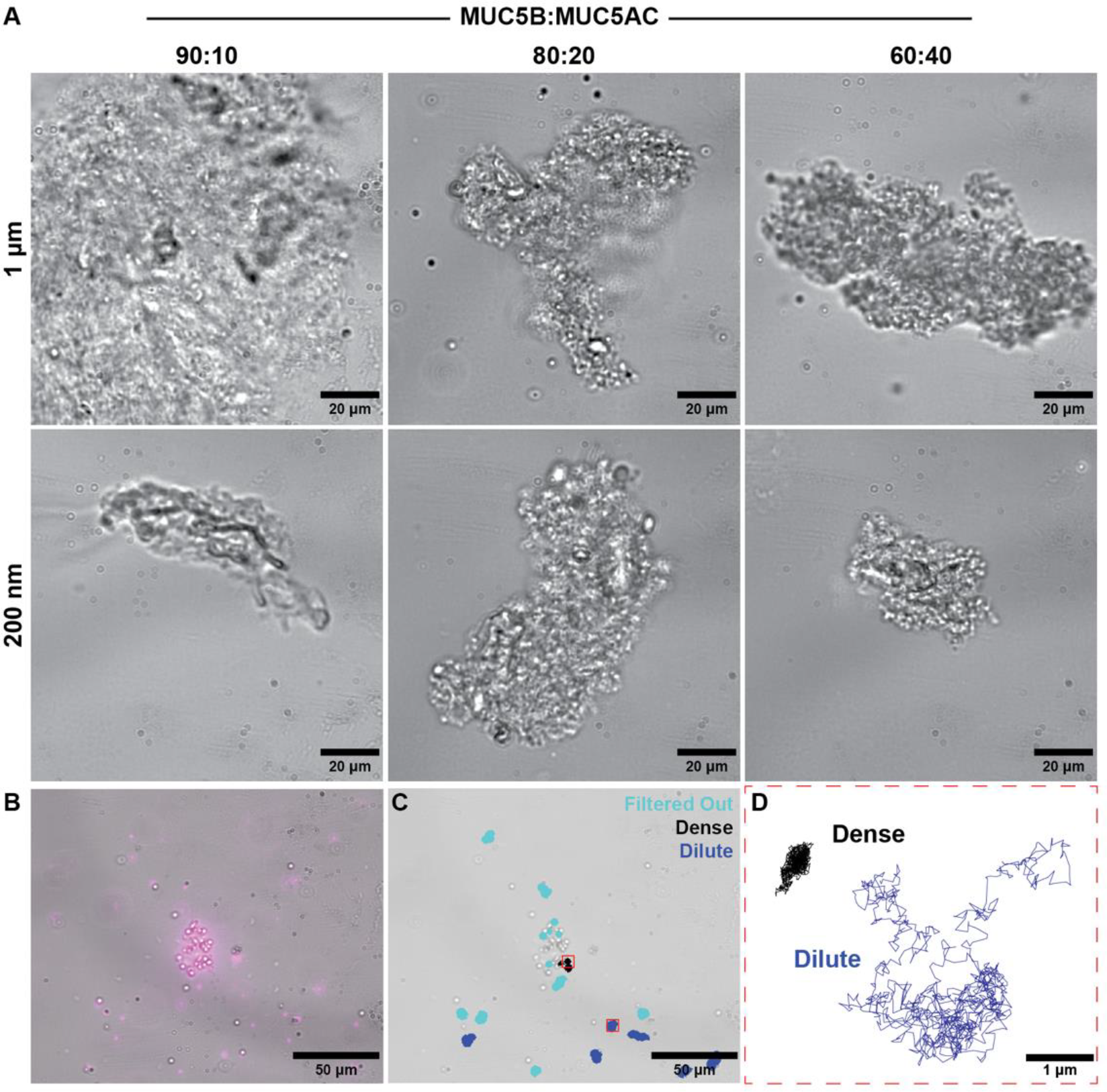
Experimental images of reconstituted mucus flakes with microbeads. (A) Brightfield images of reconstituted mucus flakes. (B) Overlay with a fluorescent image. (C) Reconstituted mucus flake overlaid with trajectories of tracked beads. (D) Higher resolution of individual trajectories of two tracked beads, one within and one outside of a flake.

Next, we explain the predictor-corrector method, then the experimental data collection, and finally we analyze data from flake-prevalent mucus samples and discuss the results.

## Particle position time series data and statistical analysis of the data

Recall Equation (1) that describes the MSD per lag time, *nτ*. For a spherical particle of radius *r* diffusing in a purely viscous medium, e.g., water or glycerol of viscosity *η*, the position time series is described by *d*-dimensional Brownian motion, for which the MSD scales linearly with all lag times:

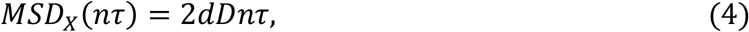

where *D* is the diffusivity of the medium given by the Stokes-Einstein relation,

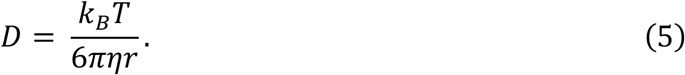

Soft biological materials like mucus are both viscous and elastic, with different responses at different frequencies of forcing. In viscoelastic materials, the MSD of passive beads is sublinear over the timescales for which the medium exhibits memory due to elastic recoil of the polymeric network. For 200 nm to 1 μm diameter passive beads in respiratory mucus, and for our microscope system, the memory timescales encompass the camera shutter timescale (1/60 sec) and the total bead observation time (30 sec). Furthermore, *over experimental timescales, each individual tracked bead approximately exhibits a sub-diffusive MSD power law*,

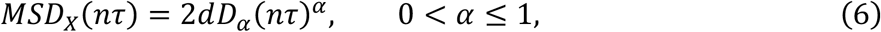

which matches that of fractional Brownian motion. It is important to emphasize that the statistical classifier for Brownian motion is one-dimensional, the diffusivity *D*, whereas *for subdiffusive fractional Brownian motion, the classifier is two-dimensional*, (*α, D*_*α*_) [15]. The power law *α* reflects the degree of sub-diffusivity of the medium surrounding each tracked particle. The pre-factor *D*_*α*_, however, has *α*-dependent units, so that one cannot compare relative numerical values across different beads, nor can one perform any numerical computations (e.g., ensemble averaging of classifiers) or cluster analysis. We return to this point momentarily. For now, we note that

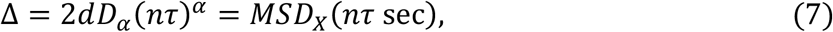

for any lag time *nτ*, e.g., *nτ* = .1 or 1 sec, has units of μm^2^ and therefore can be readily compared across multiple beads, and used for averaging or cluster analysis. Alternatively, one can also use the GSER per bead and compare the dynamic moduli *G*^′^ and *G*^′′^ of different tracked beads at any frequency, e.g., 10 or 1 sec^-1^, or perform other statistical analyses. *The potential problem with any of these single lag time or frequency projections onto a scalar quantity arises when the bead time series data has not been denoised first, and the potential for error is exaggerated for the highly sub-diffusive time series of beads inside flakes*. We illustrate this point below in Figure 3.

**Figure 2:**
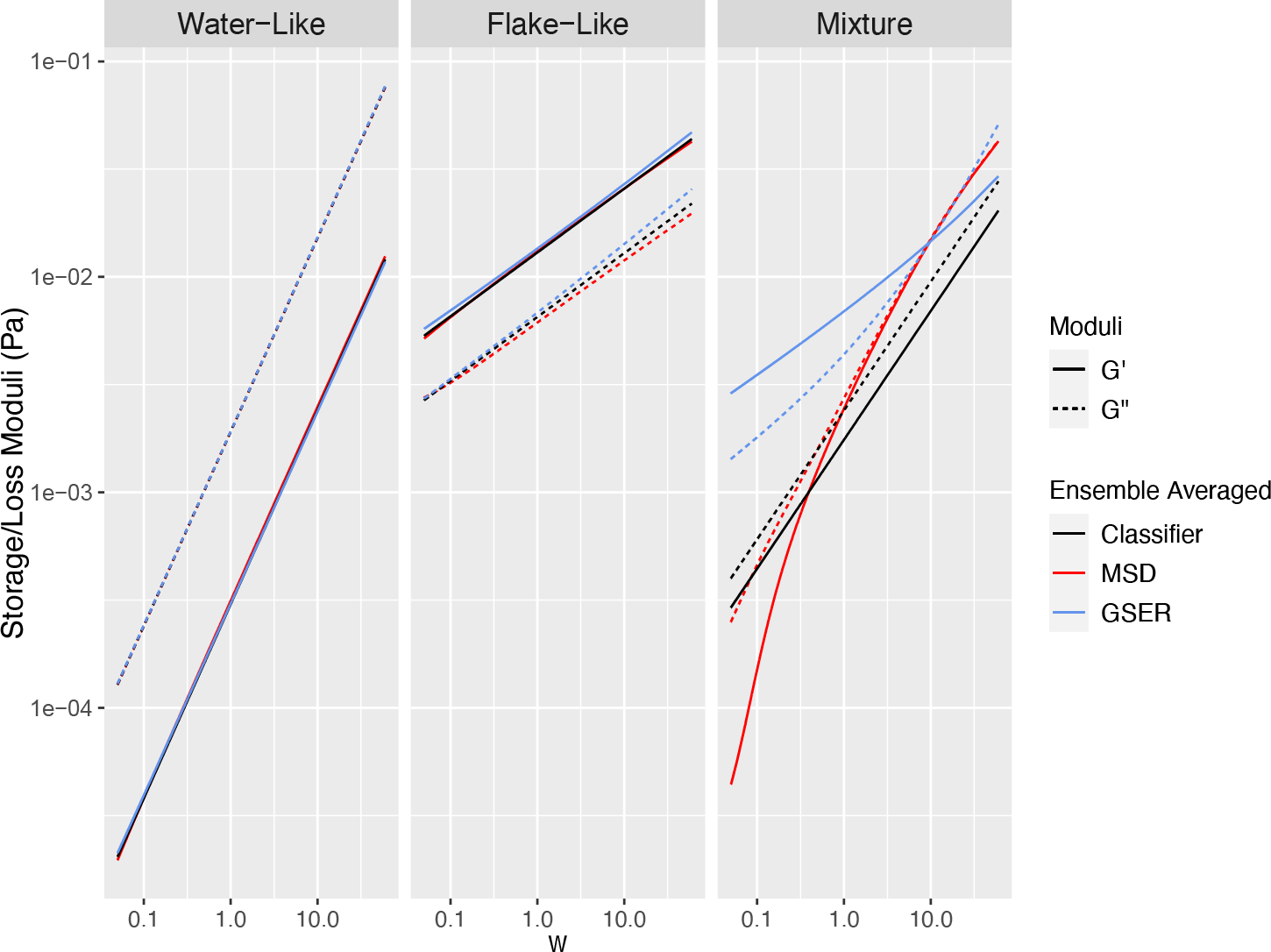
Comparison between G’ and G” computed three ways: (1) GSER applied to denoised MSD classifier (α, D_α_) per bead, eqn 2, then averaging G’ and G” in frequency space; (2) ensemble averaging over all denoised bead MSDs, then apply GSER, eqn 3 ; (3) ensemble averaging of the denoised MSD classifiers, giving a mean (α, D_α_), therefore a denoised power law ensemble-average MSD, inserted into either eqn 2 or eqn 3. These approaches are implemented for two separate homogenous clusters: a water-like cluster on the left, cluster 1, and a flake-like cluster in the middle, cluster 2, illustrating consistency in the approaches for homogeneous clusters. The mean dynamic moduli of the mixture of these two clusters are computed with each approach on the right, showing inconsistency in the approaches for heterogeneous clusters.

**Figure 3:**
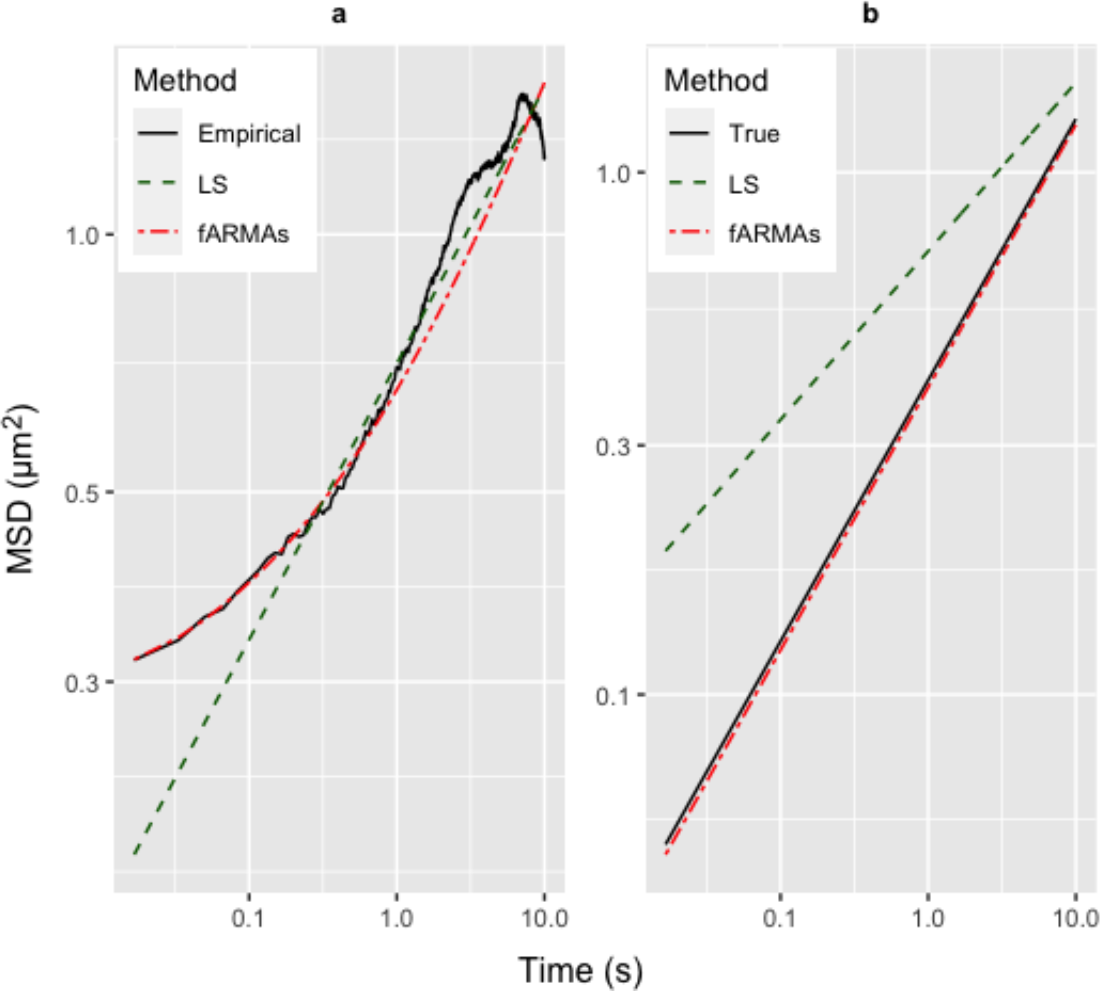
(a) Empirical and fitted MSDs to simulated time series of fBm plus noise in the low signal-to-noise range. Fitted MSDs are from LS and fARMAs. N.B. fARMAs estimates the pure fBm signal plus high and low frequency noise, which together produce empirical MSD estimates. (b) LS and denoised fARMAs pure signal MSD estimates plus the pure signal MSD.

**Figure 4:**
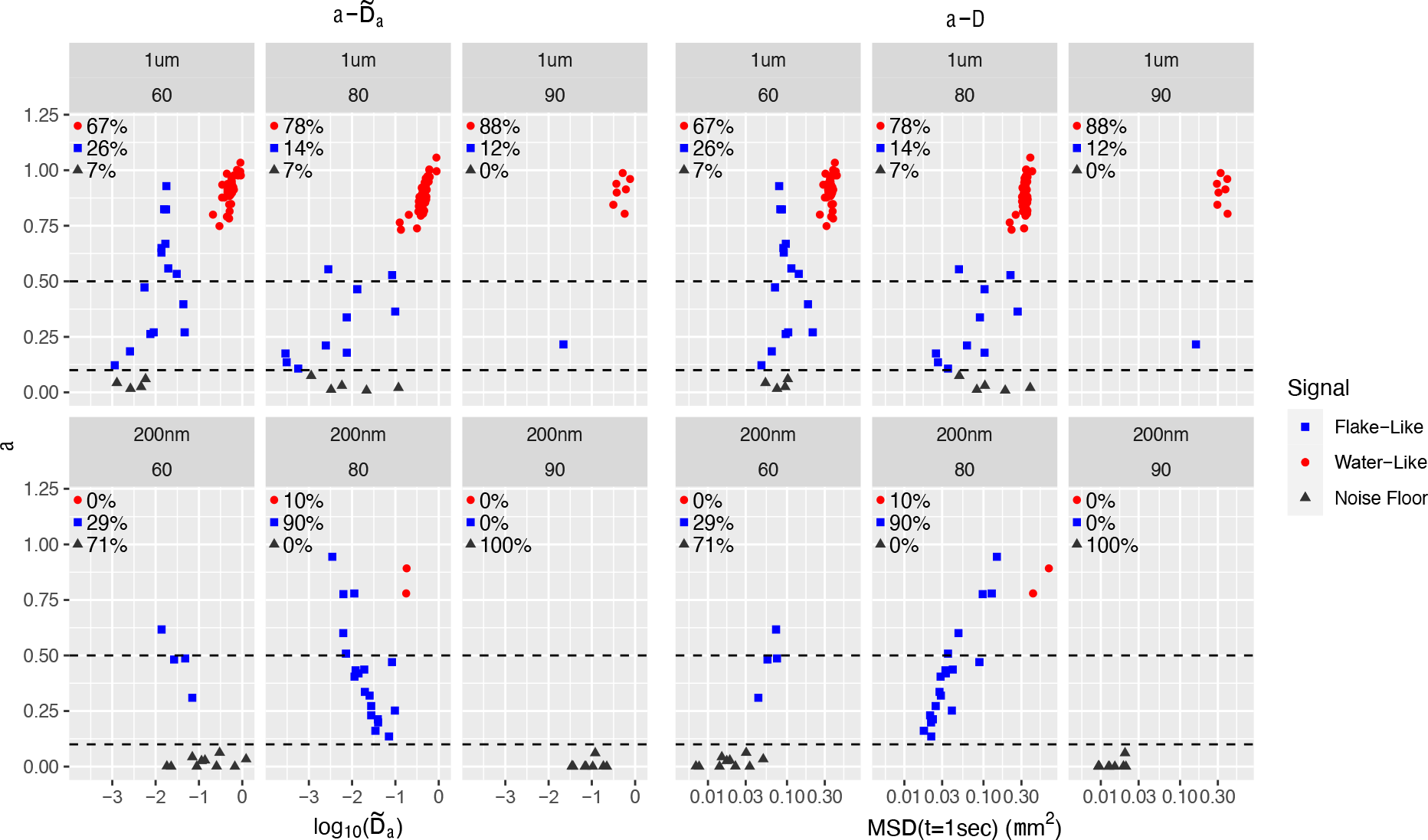
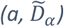 (left) and (a, Δ) (right) classifier data for individual, 1 μm (top) and 200 nm (bottom) diameter beads in three different mucin mixtures. Classifier data is visually clustered into: water-like (red), flake-like (blue), and noise floor (black) signals.

**Figure 5:**
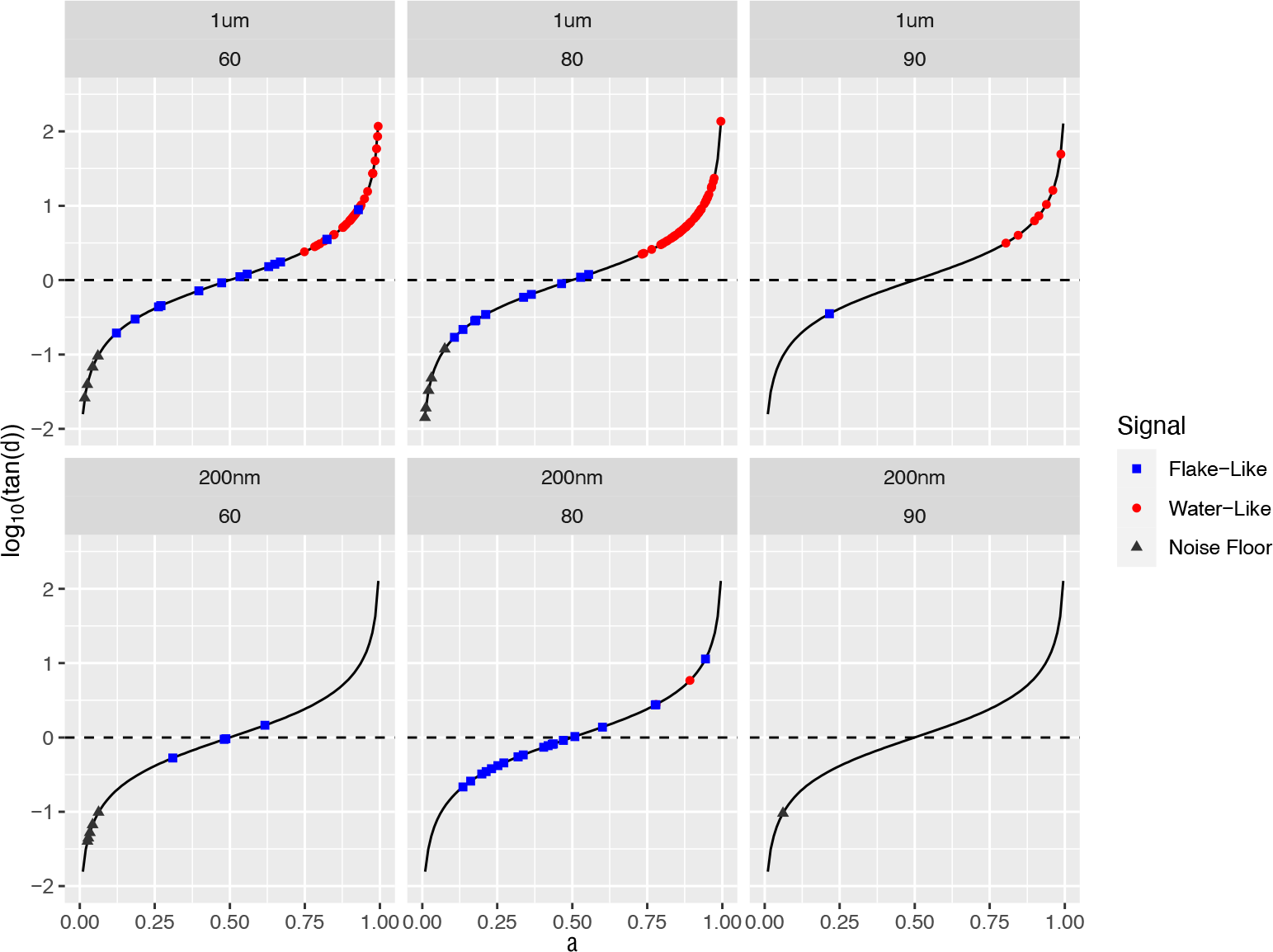
log_10_(tan(δ)) = log_10_(tan(α/2)) plots for 1μm (left) and 200 nm (right) beads in HBE + Calu3 mucus samples 60, 80, 90. Dots correspond to the scalar loss tangent metric per bead, the thin black curve is a plot of the function log_10_(tan (α/2)), the black dashed horizontal line signals the sol-gel boundary. Blue, red, black dots are from flake-like, water-like, noise floor clusters.

**Figure 6:**
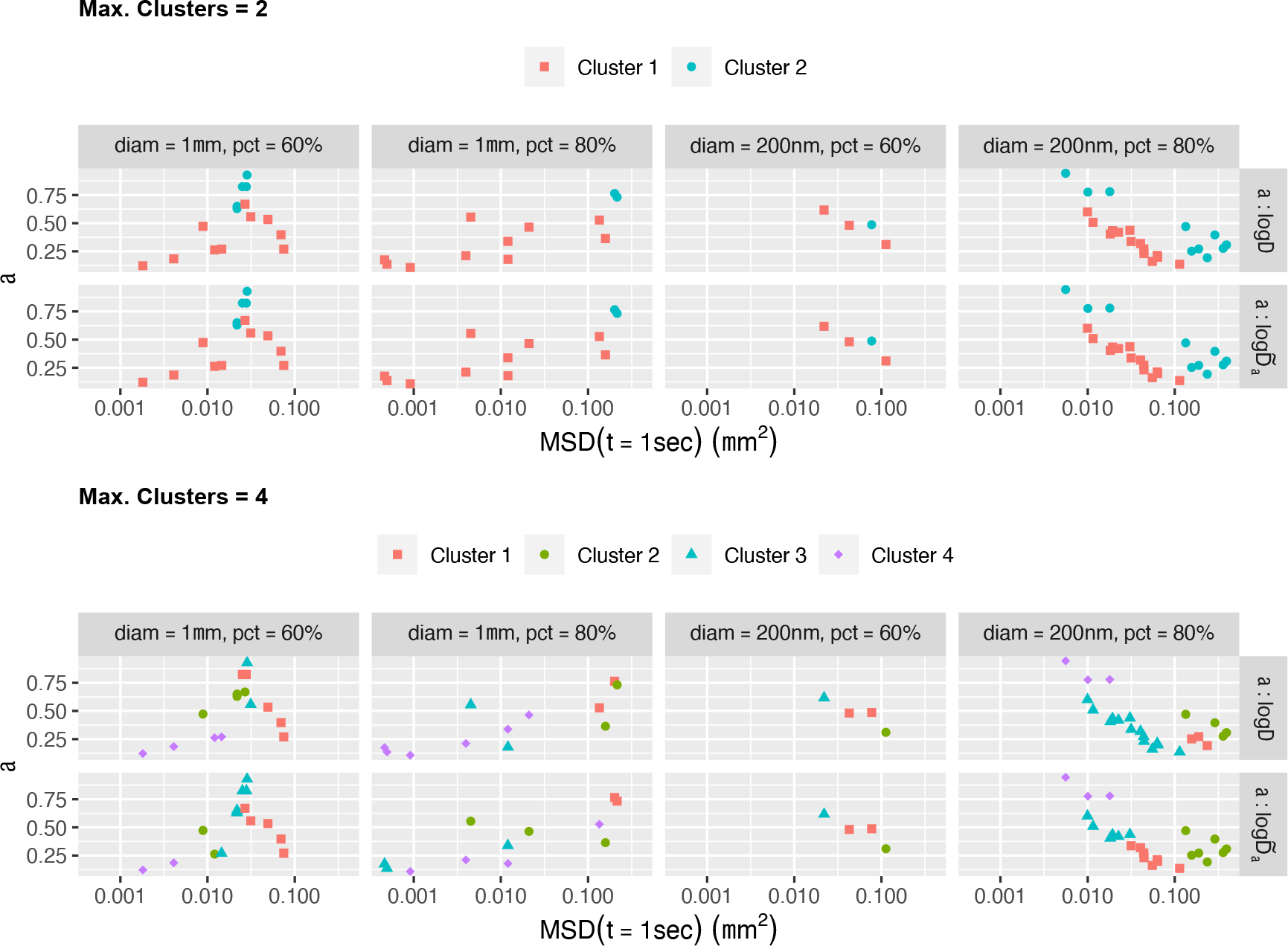
Clustering results for classifiers (α, log Δ) and (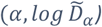 applied to all beads per sample above the noise floor with a flake-like signal, assuming a maximum of 2 clusters (top) and 4 clusters (bottom). The four samples with multiple beads are clustered using **mclust**. The clustering results for each classifier are superimposed on the same inferred data 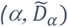, using the estimate of α on the vertical axis and the estimate of MSD(1 sec) from 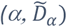 on the horizontal axis, which is independent of bead diameter, in order to compare results for each sample and bead probe.

**Figure 7:**
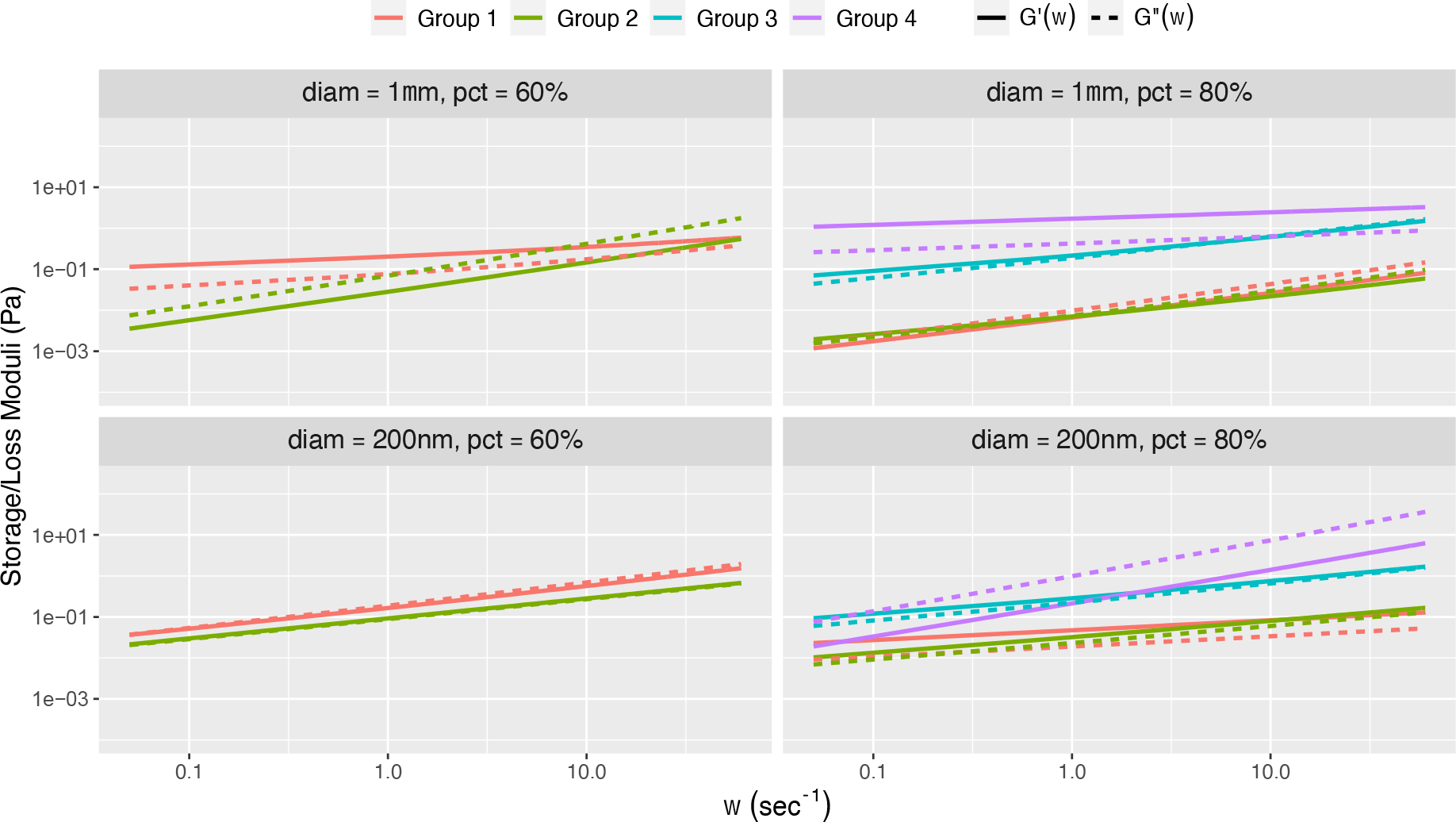
Mean storage and loss moduli for each cluster in Table 1 using the “GSER Average” data-analysis protocol. Mean + 2 s.d. intervals are included for all clusters consisting of at least two beads. Groups 1-4 are defined in Table II above.

**Figure 8:**
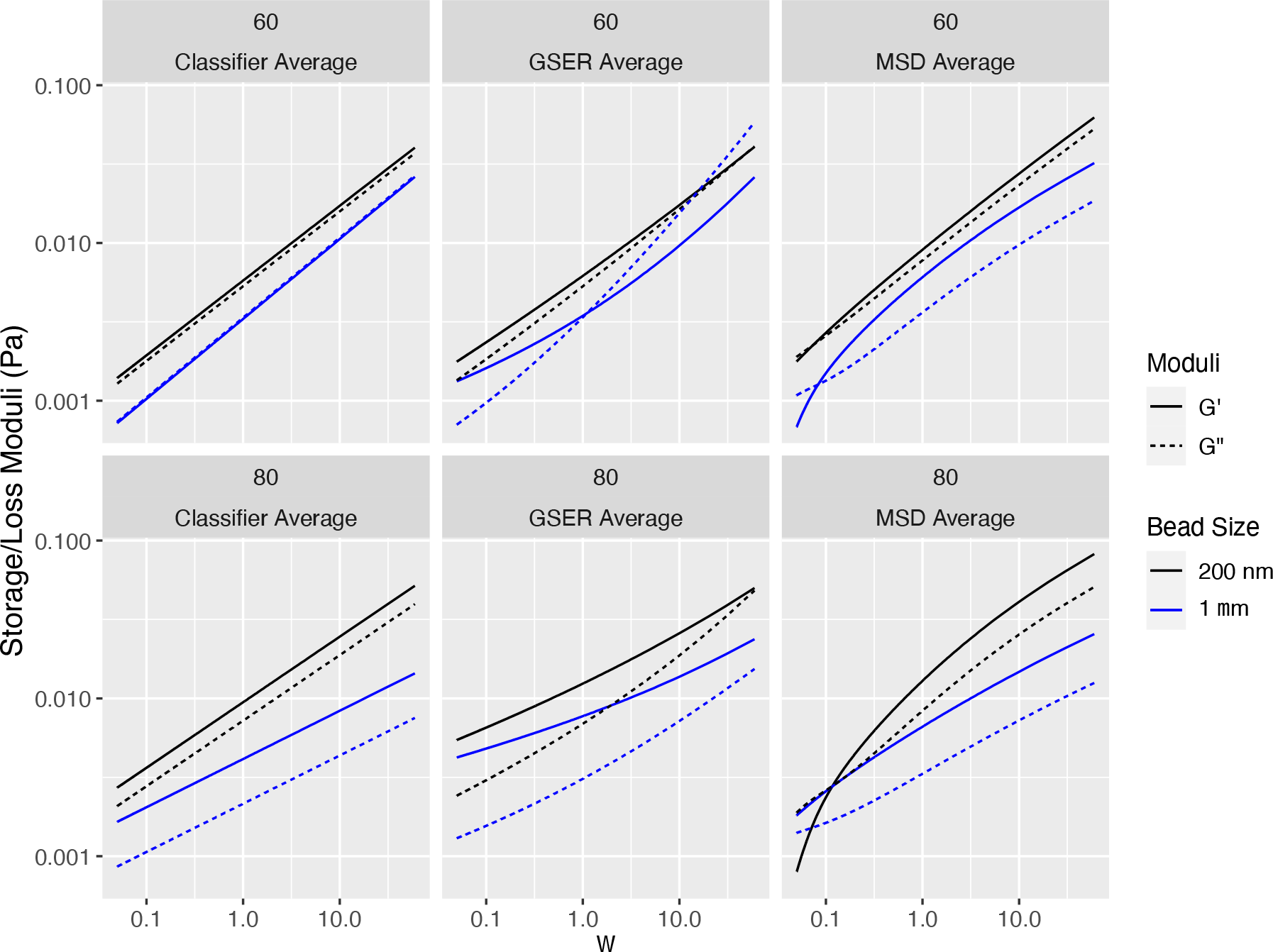
Comparison of G’ (elastic) and G” (viscous) moduli from 1 μm (blue) and 200nm (black) tracked beads in the “flake-like” cluster for the 60% (Row 1), 80% (Row 2) samples, using the Classifier Average (Col 1), the GSER Average (Col 2), and the MSD Average (Col 3). Only the GSER Average (Col 2) results are justified due to inhomogeneity of the clusters.

In our approach, *the MSD classifier* (*α, D*_*α*_) *is a denoised projection onto fractional Brownian motion (fBm)*. This classifier further provides *an analytical formula for the dynamic moduli surrounding each tracked bead* by virtue of the exact Fourier transform of any power-law MSD:

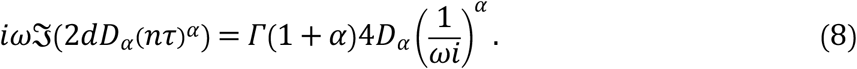

Using (8) in the GSER (2), *the denoised classifier* (*α, D*_*α*_) *per individual bead yields an exact analytical formula*, thereby avoiding numerical error:

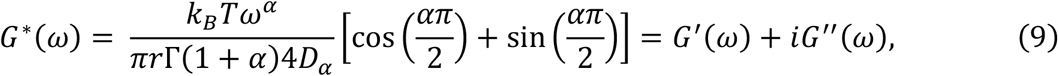

where Γ(*⋅*) is the Gamma function. From the ensemble of bead classifiers per sample per bead diameter per experiment, after appropriate rescaling of *D*_*α*_ or projection onto an appropriate scalar derived from (*α, D*_*α*_) to perform clustering, beads are filtered in a **three-stage analysis**. The **first stage** is binary: one cluster of all tracked beads with a water-like, weakly subdiffusive, signal consistent with tracking of beads in a relatively dilute mucin solution with the experimental instrumentation (thus likely *outside flakes*); a second cluster of tracked beads with a clear sub-diffusive signal bounded away from water-like signals (thus likely *within flakes*). The **second stage** is also binary, applied to the within-flake cluster. Bead increment time series are divided into those that *are, or are not, distinguishable from the noise floor of the experimental instrumentation*. The signature of the noise floor is achieved by tracking stationary beads glued to a glass plate with the experimental instrumentation, thereby identifying the classifier range where the signal is indistinguishable from the noise floor. The **third stage** involves two steps: a finer clustering of the successfully denoised outside- and within-flake clusters using a standard clustering algorithm (**mclust** [32]); followed by a homogeneity test of the clusters (Cochran’s Q test [28, 29]), the result of which guides the appropriate application of the GSER to determine equilibrium dynamic moduli of the clusters outside and within flakes. (Figure 2 below illustrates the stages of this analysis with synthetic data in the ideal scenario where clusters within and outside flakes are homogeneous.) The three-stage analysis of the experimental data thereby provides an assessment of several layers of heterogeneity in flake-burdened mucus.

Figure 2, left and middle panels, illustrate *equivalence of using forms (2) and (3) of the GSER for homogeneous clusters*. The left panel data mimics water-like signals in dilute HBE mucus and the middle panel mimics flake-like signals, both drawn from a normally distributed mean MSD classifier (*α, D*_*α*_). For the water-like signal, the means of (*α, D*_*α*_) are *μ* = (0.85, 0.35) and their standard deviations are σ = (0.03, 0.02). For the flake-like signal, we used *μ* = (0.3, 0.05) and σ = (0.07, 0.02). These clusters are not homogeneous in the strict sense of having identical (*α, D*_*α*_) values for every particle in the cluster, but rather in the broad sense of having some variation of parameters that is consistent with observed empirical data from homogeneous complex fluids. The standard deviation for *α* in the water-like signal is sufficiently small to avoid *α* > 1, which causes anomalous moduli. The two-parameter distributions were chosen independently due to lack of knowledge of correlations. The right-most panel shows the nonequivalence of the three methods for averaging of tracked bead data in a heterogeneous mixture of these two normally distributed water-like and bead-like clusters. In both homogeneous clusters all three approaches recover a power-law rheology scaling, i.e., approximately linear in log-log space, whereas only approach (3) is *guaranteed* to give power-law scaling. For the heterogeneous mixture in the right panel, methods (1) and (2) produce a non-power law scaling of *G*^′^, *G*^′′^, distinct from method (3). Neither of the methods for the heterogeneous mixture on the right-most panel has meaningful physical relevance: they average over the underlying heterogeneity without knowledge of volume fractions of flakes and dilute solution and proper weighting of each in the average. Given this additional information, assuming passive interaction between flakes and non-flake solvent, one could predict linear macrorheology of flake-burdened mucus.

## Data Analysis Methods

Our strategy is to extract the pure stochastic fluctuation signal using an extension of the *fractional autoregressive moving average* (fARMA) denoising model developed in [16]. From [33], Figure 3 below and Supplemental Material, we show that two extensions of the fARMA denoising procedure are necessary to accurately recover the pure signal in simulated truth sets consistent with the extremely low signal-to-noise (SNR) ranges of tracked beads within mucus flakes. First, fARMA is sensitive to the initial condition in the nonlinear optimization procedure. We resolve this hurdle with a least-squares predictor of the MSD followed by the fARMA corrector to ensure a stable denoising procedure. Second, the inclusion of static noise in the fARMA model, which we call fARMAs, gives more accurate estimates of the pure signal in simulated truth data with low SNR. The LS predictor – fARMAs corrector method yields a two-parameter classifier for each tracked bead of the purely entropic, medium-induced, sub-diffusive MSD over measured timescales. In Figure 3 below and more extensively in Figures 10, 11 of Supplemental Material, we illustrate the method on synthetic data, revealing estimators and standard errors across ranges of truth datasets generated from Brownian and sub-diffusive processes, with superposition of various combinations of experimental error.

**Figure 9:**
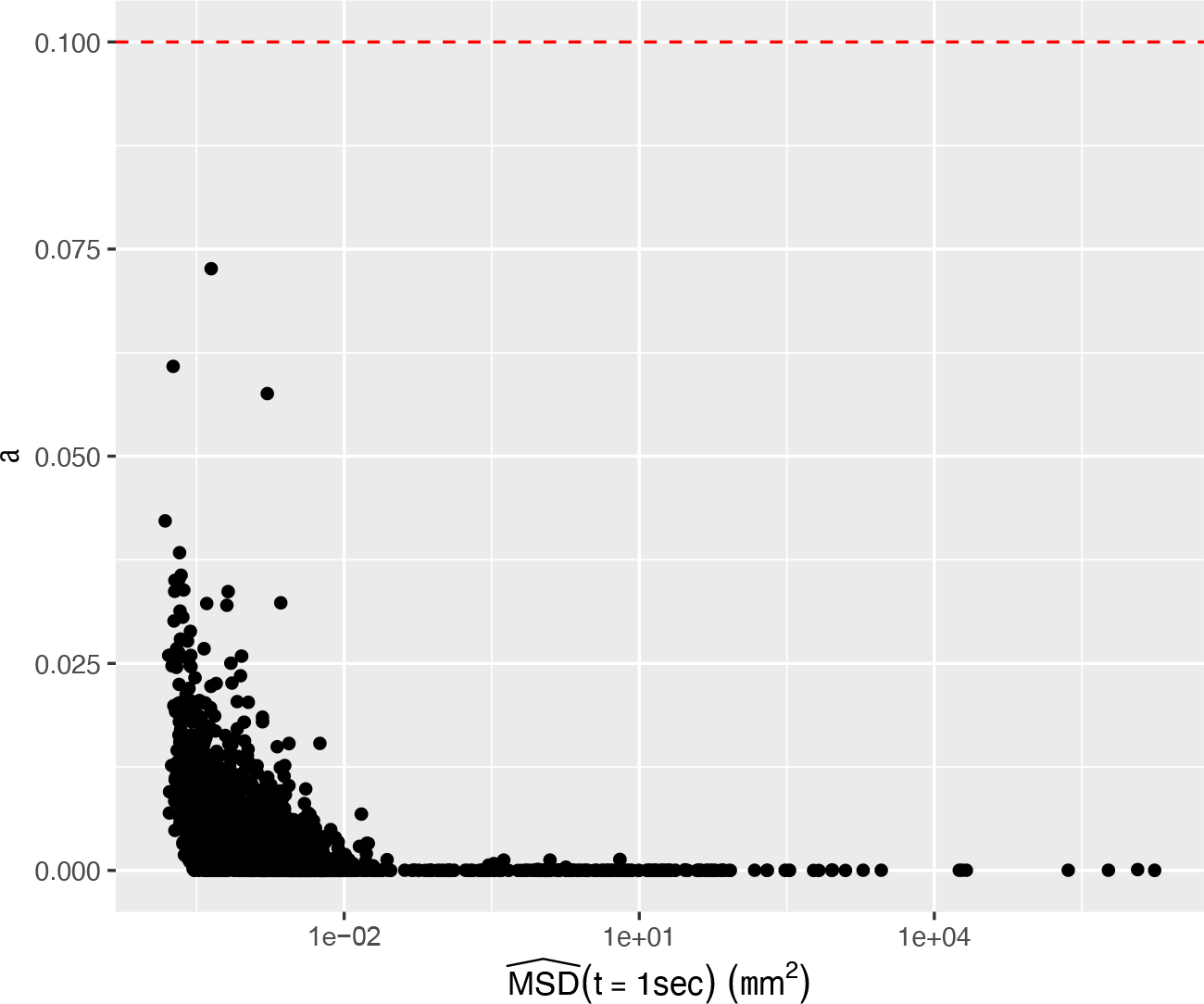
Plot of (α, Δ) values for stuck beads whose signals are purely noise. The broad range of Δ estimates, almost 10 orders of magnitude, reflect the instability in estimation of Δ when α < 0.1. Based on this data, we impose the dashed red line α = 0.1 as the noise floor cutoff.

**Figure 10:**
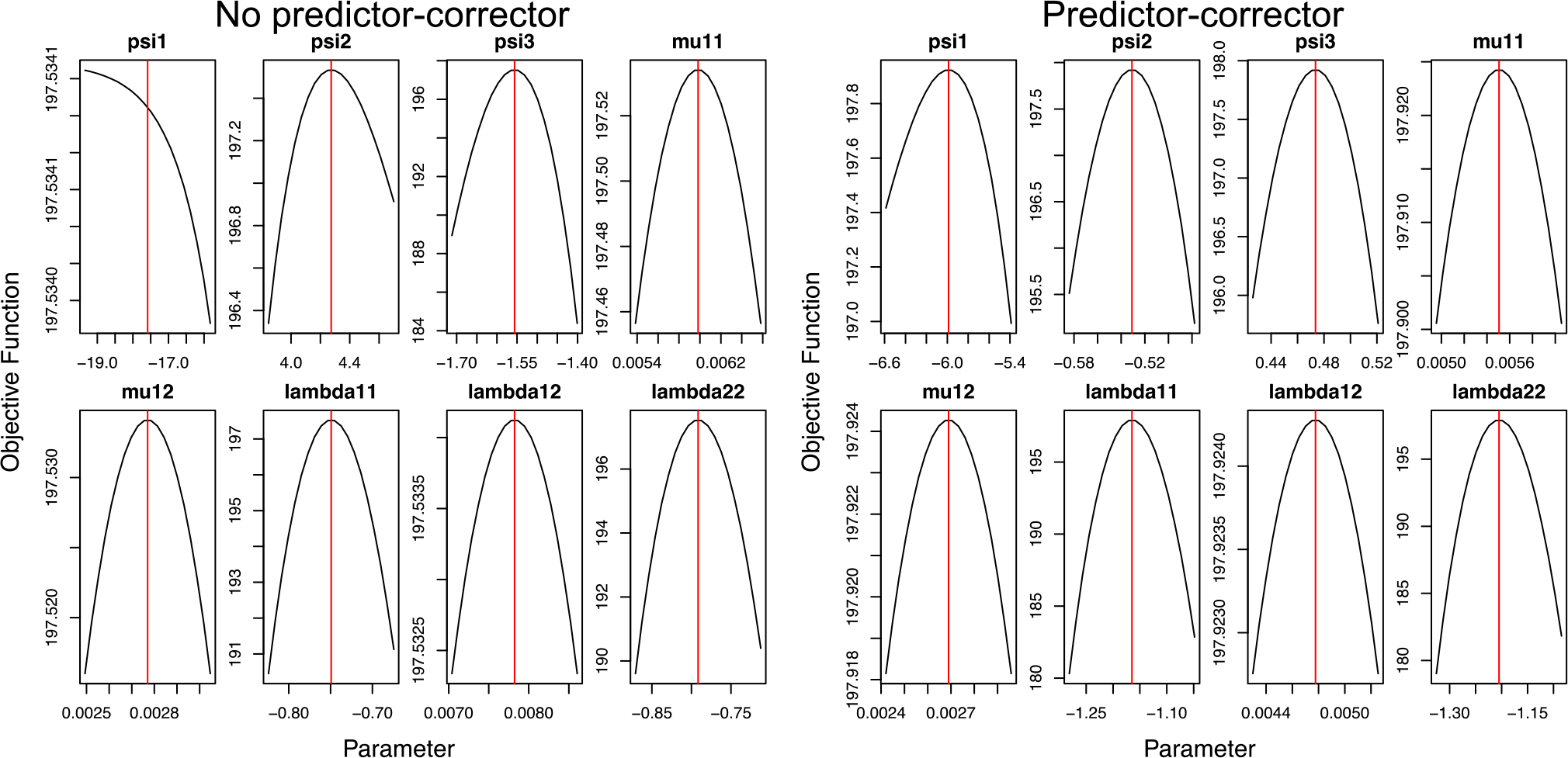
Convergence graph for optimization parameters for fARMAs. ψ_1_ *corresponds to* α, ψ_2,3_ *are noise parameters*, μ *are the linear drift parameters in x and y, and* λ *are the scaling parameters*.

**Figure 11:**
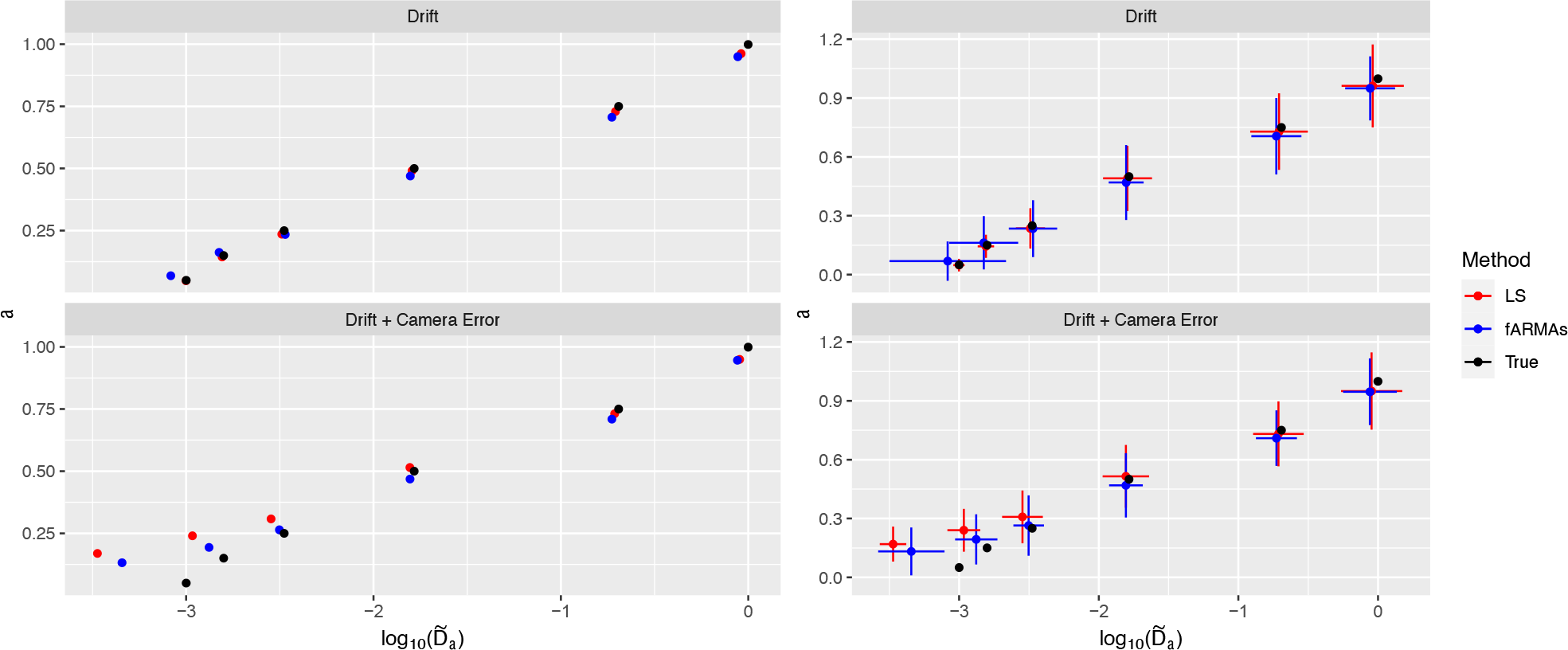
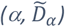 estimates for each of the six bead pairs for the two error models: low frequency, and low plus high frequency. The left panel gives the estimates from LS and LS predictor-fARMAs corrector and the true values, while the right panel superimposes crosses consisting of two times the standard error in α and 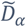.

The standard method for estimating the dimensional MSD classifier parameters (*α, D*_*α*_) is via least-squares (LS), which consists of fitting the slope and intercept in the relation [31]

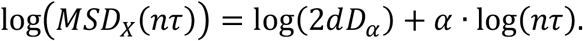

Since the empirical MSD is contaminated by noise at both the smallest (static and dynamic camera error) [27] and largest (drift of the sample) timescales [9,10], lag times *nτ* in a restricted bandwidth are often used. This approach does not remove the noise from the selected bandwidth: it merely assumes that noise contamination is negligible therein. However, this assumption is violated precisely in cases of interest here, for dense flakes, where the particle mobility is within or near the noise floor. To illustrate, Figure 3a displays the MSD of fBm-contaminated by a noise floor of magnitude σ^2^, for which the MSD is given by

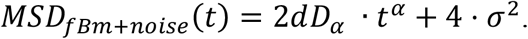

LS is fit to the bandwidth of 0.1-2 sec, and we see that the line of best fit agrees very well with that of the empirical MSD over the bandwidth. However, this fit is to the MSD consisting of signal plus noise: Figure 3b shows that the LS fit is very different from the MSD of the pure signal. In contrast, we employ an extension of the fARMA denoising model of [15] that includes static noise, denoted fARMAs [33], using the LS fit as a predictor step, which accurately recovers the pure signal in Figure 3b. Details of the fARMAs model and associated fitting procedure – including our predictor-corrector method – are given in Supplemental Material.

From here on, we use the LS (predictor) -- fARMAs (corrector) procedure to estimate the diffusion parameters. The resulting *fractional-diffusion MSD classifier*, (*α, D*_*α*_), is estimated per tracked bead, then converted to have comparable units (using (7) or Equation (10) below) for quantitative analysis, in particular homogeneity testing and clustering.

## Clustering Analysis

Let 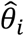denote the estimate for bead trajectory *i* of its true classifier value *θ*_*i*_. A simple choice of classifier is *θ* = (*α*, log Δ), where Δ = *MSD*_*fBm*_(*t* = 1 *sec*) via (7), with an in-depth discussion of various classifiers to follow. Given the classifiers of *M* trajectories, clustering is performed using the R package **mclust** [32] using finite normal mixture models. That is, the classifier estimates 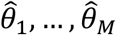 are clustered, in the most general case, according to the multivariate normal mixture model

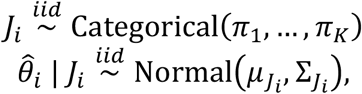

where *K* is the number of clusters and *J*_*i*_ *∈* {1, …, *K*} is the cluster to which particle *i* belongs. **mclust** estimates the posterior membership probability

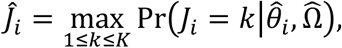

where 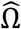 is the maximum likelihood estimate (MLE) of the clustering model parameters Ω = {(*π*_*k*_, *μ*_*k*_, Σ_*k*_): *k* = 1, …, *K*}. **mclust** chooses *K* by fitting all models with *K ∈* {1, …, *K*_*max*_} for user-specified *K*_*max*_, and selects the one with the lowest value of the Bayesian information criterion (BIC). It can also impose various constraints on the variance matrices, e.g., proportional variances Σ_*k*_ = *τ*_*k*_ · Σ_0_ or diagonal variances 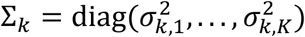 (there are 14 possible variance restrictions in total, see Table 3 of [32]). This is particularly useful when some of the clusters are expected to consist of only a handful of particles, in which case the corresponding unconstrained variance matrices can be very poorly estimated. Once again, **mclust** estimates the posterior membership probabilities 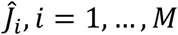 from the best-fitting model (in terms of BIC) among all combinations of cluster sizes *K* and variance matrix constraints.

## Homogeneity Testing

Now suppose that in addition to 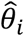, for each trajectory *i* we have an estimate of the variance of its classifier, 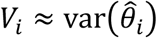. Such a variance estimator is obtained as a direct by-product of the fARMA denoising procedure. In order to test whether the 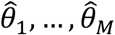 are obtained from a homogeneous cluster, let 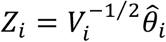 and 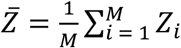 . Then under the null hypothesis of homogeneity

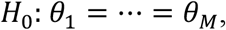

The so-called Cochran’s Q statistic [29,29]

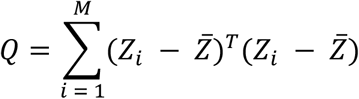

has a 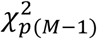 distribution under *H*_0_, where *p* is the common number of elements of each 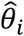 in our case we have *p* = 2. The assumption of homogeneity is then rejected at the ε level when *Q* > *C*_*∈*_, and *C*_*∈*_ is such that 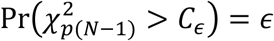.

## Tracked Particle Classifiers

A natural choice for the classifier of sub-diffusive particle trajectories is *θ* = (*α*, log*D*_*α*_), where use of log*D*_*α*_ reflects the empirical finding that *D*_*α*_ tends to vary by an order of magnitude within a given experimental setting, i.e., errors in fitting the slope extrapolate to fitting the y-intercept. However, the units of *D*_*α*_ are *α*-dependent, which precludes direct comparisons across beads. There are multiple ways to address this, e.g., one can simply evaluate the MSD at a chosen lag time *nτ*, recall the discussion surrounding equation (7) above, which gives units of μm^2^ for all beads, and thus admits inter-bead comparisons. A downside of this classifier is that the choice of MSD lag-time *nτ* is arbitrary, and one could have chosen any *nτ* to evaluate the MSD. Any clustering based on these arbitrary choices would have to be tested for robustness of the chosen timescale. We note, however, that if the ensemble data (*α*, log*D*_*α*_) has been estimated by the LS-fARMA method, then it has already used all of the denoised experimental data to produce the denoised classifier (*α, D*_*α*_) for each bead, which produces a denoised power-law MSD estimate, and therefore the clustering outcome will be relatively robust for all choices of MSD (*nτ*). This is not the case for standard LS-estimates of (*α, D*_*α*_), illustrated in Figure 3; namely, the LS estimates of MSD have different errors from the true signal MSD at every lag-time *nτ*! An alternative approach to evaluation of MSD at lag-time *nτ* for some *n* is to non-dimensionalize *D*_*α*_, labelled 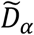. Once again, we emphasize that we use the LS-fARMAs method on the experimental data to produce the denoised classifier (*α, D*_*α*_) for each bead. Using only experimental scales and a reference fluid relevant to mucus, *we define a power law α*-, *spatial dimension d*-, *and bead radius r-dependent rescaling of D*_*α*_ normalized with respect to the diffusivity of a bead of the same radius in water,

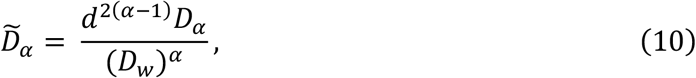

where *D*_*w*_is the viscous diffusivity of water for particles of radius *r*. Note that

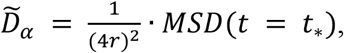

where *t*_*∗*_ = (2*r*)^2^⁄*D*_*w*_. In other words, 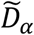 can be interpreted as a multiple of the MSD at a time *t*_*∗*_which depends on the particle radius *r*, instead of an experimental lag time *nτ*. For 1 μm beads, *t*_*∗*_ = 2.33 *sec*, in the middle of the experimental lag times, and for 200 nm beads *t*_*∗*_ = 0.018 *sec*, just above the minimum experimental lag time *τ*. Of note, the homogeneity test produces identical results when applied to (*α*, logΔ) or 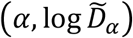 since they are linear transformations of each other. The same is true for the **mclust** clustering algorithm with unconstrained variances.

For viscoelastic media, a fundamental property is that, instead of a scalar metric (viscosity for purely viscous, elasticity for purely solid, materials) the viscous and elastic moduli are functions of frequency. In this respect, a standard viscoelasticity metric is the loss tangent:

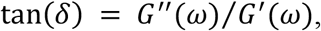

which in general is frequency-dependent. When *δ* > *π/*4: tan(*δ*) > 1 *and the material is sol-like* (i.e., loss or viscosity dominated), whereas when *δ* < *π/*4: tan(*δ*) < 1 *and the material is gel-like* (i.e., storage or elasticity dominated).

*For pure power-law viscoelastic materials*, tan(*δ*) *collapses to a scalar function of the MSD power law exponent* α, *independent of both frequency w and the MSD pre-factor D*_α_. From the GSER (2) and the power-law moduli formulae (9):

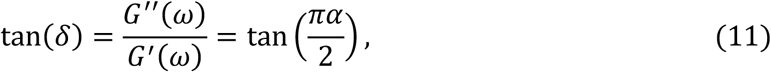

such that 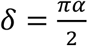. Thus, *gel-like phases correspond to* 0 < *α* < .5 and *sol-like phases correspond to* . 5 < *α* ≤ 1. For easier graphical visibility, we plot log_10_(tan *δ*), shifting the gel/sol cutoff in log_10_(tan *δ*) to 0, so that *positive values of* log_10_(tan *δ*) *are sol-like and negative values gel-like*. All water-like data is sol-like, all noise-floor data is gel-like, and all flake-like data lies between these extremes. We are interested especially in the percentage of sol-like and gel-like signals in flake-like data as a coarse scalar metric of heterogeneity.

## Materials and Data Collection

1 μm and 200 nm beads are embedded in a mixture of purified mucins (MUC5B, MUC5AC) derived from either HBE (predominantly MUC5B) or the human airway epithelial Calu3 cell line (MUC5AC). We chose to study the effect of the changed ratio of muc5b to muc5ac during the progression of CF [36]. Three different combinations of mucin are used: 90%:10% (non-CF), 80%:20% (mild CF), and 60%:40% (severe CF) HBE:Calu3-MUC5AC. 2% mucin solution was used for all conditions.

The flakes were made by mixing beads into the mucin solution followed by injecting that mixture into a calcium chloride solution. Imaging revealed both within-flake and outside-flake beads. There are two reasons that some beads are not found in flakes at the time of tracking: 1) the process doesn’t have 100% encapsulation, and 2) beads formerly inside may diffuse out.

Before diffusion parameters are estimated using fARMAs, first all beads are passed through a series of filters; the most notable of which is the proximity filter which ensures that the only forces acting on the bead are from the medium itself and not from other beads in proximity. A full analysis of the data and details on the pre-filtering is provided in the Supplementary Information.

## Results on Experimental Datasets

### Coarse Clustering: Beads Within vs. Outside Flakes

Figure 4 gives the two-parameter classifier estimates for 1 μm and 200 nm diameter beads in 3 mucus mixtures: HBE + Calu3 60, 80 and 90 as described in the **Materials and Data Collection**. Two different scales are used for comparison: 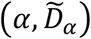and (*α*, Δ = *MSD*_*X*_(*t* = 1 *sec*)). Recall that after taking logarithms of 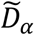 and Δ, the two scales are linear transformations of each other. An advantage of using Δ = *MSD*_*X*_(*t* = 1 *sec*) is that it allows for comparison between 1 μm and 200 nm beads, which cannot be done with 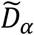 since it is bead radius-dependent. *All datasets are visually divided into three clusters: water-like or outside flakes, within flakes but entangled in the noise floor, and flake-like with recoverable signal*. The designation of tracked beads are indistinguishable from the “noise floor” is based on a stuck bead analysis provided in the Supplementary Material with cutoff *α* < 0.1. The flake-like data with recoverable pure signal exhibits significantly sub-diffusive behavior yet above the noise floor. In [33], this classifier-based coarse clustering is shown, by comparison with visual data from all experimental datasets as in Figure 1, to have extremely high accuracy in predicting beads within or at the periphery of flakes versus outside flakes.

We draw further consequences from the comparison of 1 μm and 200 nm diameter probes in all three samples.

1. ∼ 74% of 1 μm diameter beads exhibit water-like signals versus ∼ 5% of 200 nm diameter beads (0/23 in the 60 and 90 samples and 2/21 in the 80 sample). These statistics of entry into flakes far outweigh, based on normal diffusion, the 5:1 expected flake encounter frequency of 200 nm vs. 1 μm diameter beads. (The encounter frequency ratio for sub-diffusive beads is not known, to our knowledge.) *The results support several flake structure properties:* (i) flake encounters of 200 nm beads almost always lead to entry, implying *a high percentage of pores at the boundary of flakes is above 200 nm*; (ii) flake encounters of 1 μm diameter beads have a *high (*∼*74%) failure rate of entry*, even though each freely diffusing bead likely has many flake encounters. Assuming purely steric interactions, this data suggests a maximum 26% of boundary pores of flakes are above 1 μm, and a minimum 74% of boundary pores of flakes are below 1 μm, with a very low % of boundary pores below 200 nm. A series of steps is required to do better than these crude estimates of pore-size distributions. *If*: (i) the interactions of beads with mucus are purely steric; (ii) the volume % of flakes versus dilute solution is known; (iii) the relative sub-diffusive bead mobilities within flakes and dilute solution can be transformed to flake encounter frequencies; and (iv) the pore-size distributions at the boundary of flakes is known; *then* it would be possible to estimate the quasi-stationary percentages of 200 nm and 1 μm diameter beads in flakes and solvent. An interesting open challenge is: given the experimental data on percentages of 200 nm, 500 nm, 1 μm diameter beads in flakes and solvent, infer the pore-size distributions across the entire sample. This inverse problem is solvable for normally diffusive probes, cf. [33], but not for sub-diffusive probes, which we have shown here is the case down to 200 nm diameters in mucus flakes and in remaining dilute solution.
2. The total within-flake data gives further insights into internal flake pore morphology. The *200 nm data* across the three different samples suggests *a non-monotone pore structure transition* from the 60 to 80 to 90 samples: in the 60 sample, 10/14 *beads enter flakes and become trapped with SNR buried in the noise floor*; in the 80 sample, 0 *beads enter a flake and become trapped with SNR in the noise floor*; in the 90 sample, *all 9 beads enter and become trapped with SNR indistinguishable from the noise floor*. Figure 3 reflects a nonmonotonicity in rheology since it is deduced from the same data. This non-monotonicity could be a result of small sample size, or the stochastic nature of mucin polymer phase separation in the formation of flakes. Additional dedicated experiments coupled with advances in analytical methods in the previous comment are necessary.
3. The *1* μm *bead data for 60, 80 samples* are consistent with a *broad pore size distribution* so that beads randomly sample outside and within flakes, with *some within-flake beads in pores larger than 1* μm *and others entering and being trapped in pores slightly larger than a micron, the latter reflected by beads in the noise floor*. The *1* μm, *90 sample dataset is smaller, yet suggestive that the pore-size distribution is broader with many multi-micron pores, so that beads freely enter and escape flakes. This would be consistent with a less dense flake formation at the smallest, 10%, Calu3 MUC5AC consistent with early CF progression*.
4. Use of tan *δ* = 1, equivalently *α* = 0.5, as the sol-gel boundary is a fairly accurate predictor for water-like vs. flake-like classification of tracked beads. It agrees with our two-parameter classifier for six of the eight samples. As could be expected in a phase separation process, two samples have overlapping *α* values above 0.5, and their cluster separation is only captured by using the two-parameter classifier as evidenced in Figure 6 below.

Figure *5* gives a visual equivalent of the *α* = 0.5 cutoff in Figure 4 as the separation between sol-like and gel-like local sample properties surrounding each bead. Formula (11) shows that using *α* as a scalar metric is equivalent to the loss tangent for power law fluids. Furthermore, our predictor-corrector method to infer (*α, D*_*α*_) is extremely robust in estimation of *α*. Using only *α* as a classifier does a remarkably good job of predicting the clusters of beads within each sample, whereas *D*_*α*_ (upon converting to comparable units Δ or 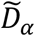) basically plays the role of a corrector in singling out the few flake-like beads with values of *α* that could have been classified as water-like. This coarse scalar metric does not, of course, convey the dynamic moduli surrounding each bead, and the reason to estimate (*α, D*_*α*_) as accurately as possible. *We now proceed to two-parameter clustering and estimates of dynamic moduli, focusing on within-flake beads at 200 nm and 1* μm *diameter scales and tracking data distinguishable from the noise floor*.

## Clustering Analysis

Figure *6* displays the algorithmic *clustering results* obtained from **mclust** for *beads above the noise floor* (*α* > 0.1) *and with a flake-like signal* for each of the two-dimensional classifiers: (i) (*α*, logΔ) and (ii) (*α*, log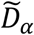). In both cases the plotting axes are *α* vs. logΔ.

The top panel of Figure 6 presents results for which the maximum number of clusters is set to *K*_*max*_ = 2, where both (*α*, log_10_ Δ) and 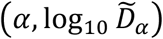 classifiers return identical clustering. The bottom panel of Figure 6 presents results for which the maximum number of clusters is set to *K*_*max*_ = 4, surmising there may be more than one or two distinguishable signals for flake-like beads.

The experiments with 1*μm* diameter beads in 90% concentration and 200nm diameter beads in 60% concentration contain too few beads above the noise floor to draw meaningful conclusions. The following observations apply to the remaining experiments.

Overall, the two classifiers, (*α*, log_10_ Δ) and 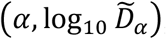, give similar results for distinguishing highest and lowest mobility clusters, but differ in how they cluster the intermediate range data. This is because **mclust** has selected a constrained variance matrix, which is typically not invariant to the linear transformation between (*α*, log_10_ Δ) and 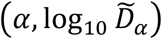.

The BIC-based selection of the number of clusters employed by **mclust** picks more than two clusters for all four experiments. However, the **mclust** mixture-normal clustering approach struggles with non-elliptical clusters. That is, each of the *K* **mclust** clusters can be identified with an ellipse, and the data points belonging to it are those for which the “natural” distance to the ellipse centroid -- i.e., along its principal axes and inversely proportional to the length of these axes -- is smaller than for any other cluster’s ellipse. This is why, e.g., the results with *K*_*max*_ = 2 for the 1 *μm* beads at 60% concentration may seem unexpected: while a visual clustering might separate the data points between left and right, **mclust** separates them between top and bottom, because the small elliptical cluster at the top and large elliptical cluster at the bottom resulted in the smallest within-ellipse natural distance between data points and ellipse centroids.

The extent to which the visual clusters are not elliptical also explains why **mclust** typically selects more than two clusters, i.e., preferring to split highly non-elliptical clusters into several more elliptical ones. That being said, clusters corresponding to perfectly homogeneous particles according to the setup in the Homogeneity Testing section are indeed perfectly elliptical.

Taking all visual and algorithmic clustering results into account, we select a single clustering per experiment from the two panels of Figure *6* summarized in Table 1.

**Table 1:**
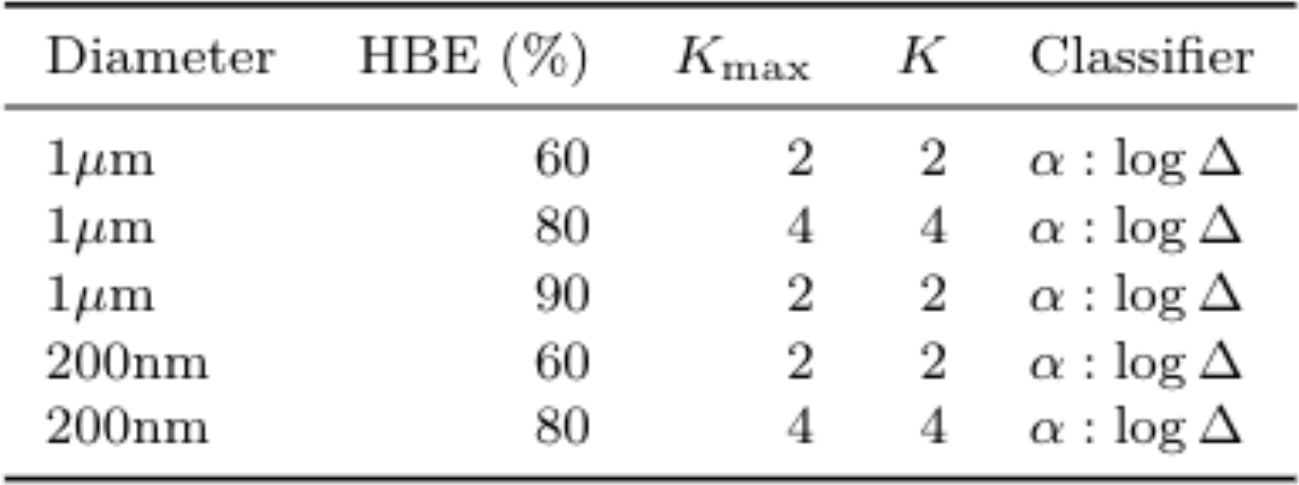
Selected cluster assignment based on visual clustering and the algorithmic clustering results of Figure 6. *K*_*max*_ *refers to the max number of clusters and K refers to the optimal number of clusters*

| Diameter | HBE (%) | $K_{max}$ | $K$ | Classifier |
| --- | --- | --- | --- | --- |
| $1\ \mu\text{m}$ | 60 | 2 | 2 | $\alpha : \log \Delta$ |
| $1\ \mu\text{m}$ | 80 | 4 | 4 | $\alpha : \log \Delta$ |
| $1\ \mu\text{m}$ | 90 | 2 | 2 | $\alpha : \log \Delta$ |
| 200nm | 60 | 2 | 2 | $\alpha : \log \Delta$ |
| 200nm | 80 | 4 | 4 | $\alpha : \log \Delta$ |

It should be noted that while (*α*, logΔ) is the chosen classifier for each of the five samples, for 4 out of the five samples the 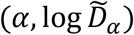 classifier returns the same clusters.

## Homogeneity Testing within Clusters

In order to assess within-cluster homogeneity, Table 2 presents the results of Cochran’s Q test applied to each of the clusters of each experiment described in Table 1, along with the cluster of beads with a water-like signal.

**Table 2:**
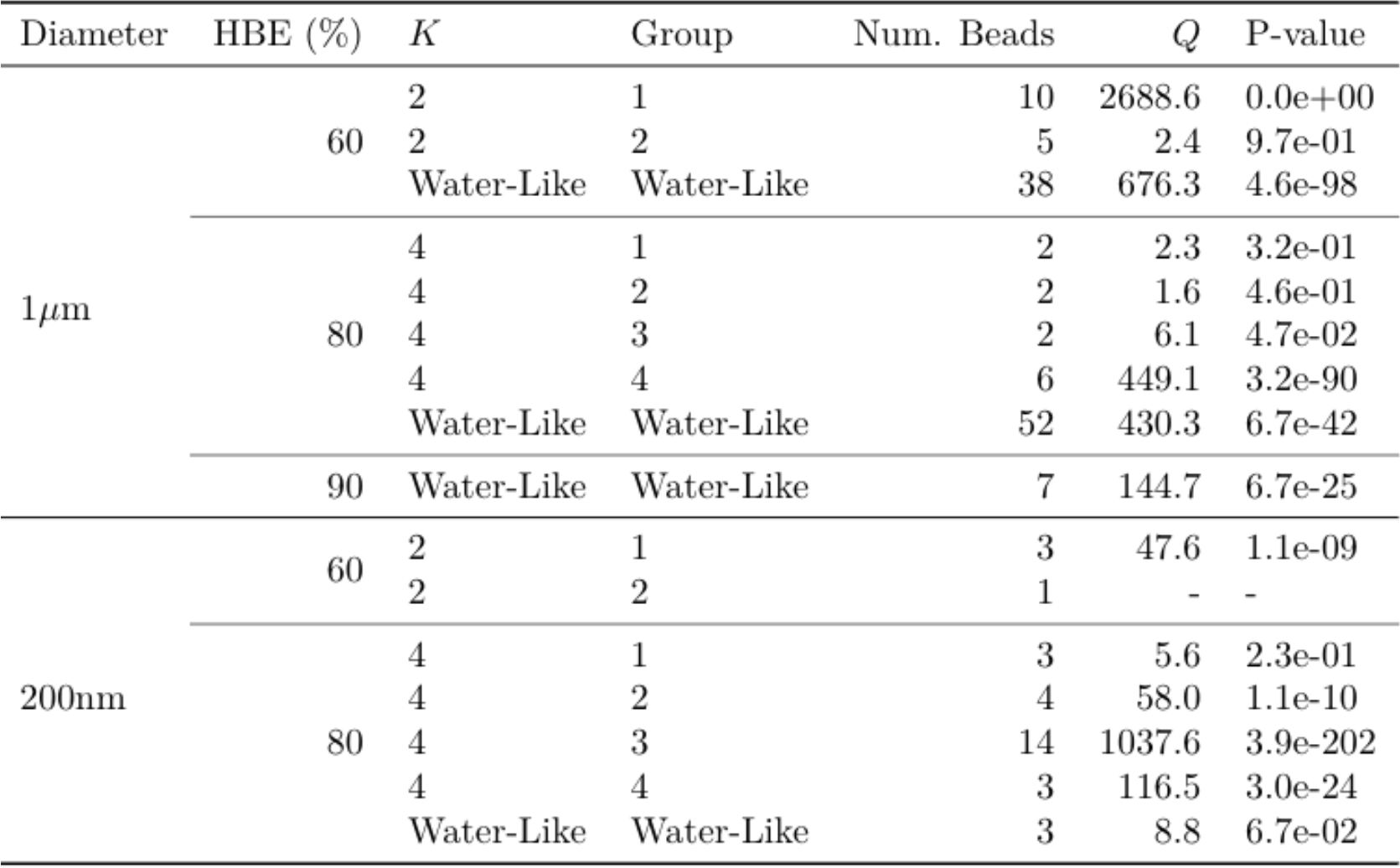
Cochran’s homogeneity test results for the clusters reported in Table *I*

All but five *p-values are extremely small*, so that Cochran’s Q test *strongly rejects cluster homogeneity*. The lack of homogeneity is expected for flake-like clusters as the data comes from different flakes, each with pore structures arising from the stochastic phase separation process of flake formation. Furthermore, only one water-like / outside-flake cluster passes a homogeneity test, and this is a weak conclusion since it is based on three beads. The outside-flake heterogeneity is also consistent with the stochastic dilution outside of phase-separated flakes, creating dilute phases that vary with relative proximity to flakes and degrees of dilution, with signals ranging from pure water-like to varying mucin concentrations.

This determination of heterogeneity, both between and within the coarse within-flake and outside-flake bead signals, strongly supports the following **data-analysis protocol**: (1) for each batch sample, separately for sub-samples with 200 nm and 1 μm beads, use coarse clustering on the (*α*, logΔ) or 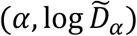 classifier per bead to cluster beads within and outside of flakes; (2) when (here, both) clusters fail the homogeneity test, further apply clustering to the within-flake and outside-flake ensembles; (3) for each cluster within and outside flakes, apply the GSER with the denoised MSD for each bead; (4) for homogeneous clusters, either of the three methods illustrated in Figure 2 can be used, while for heterogeneous clusters, *average the dynamic moduli from all beads within each cluster*; (5) assess cluster to cluster heterogeneity at the coarse scale of the within and outside flake ensembles, and then at the finer scale of clusters within flakes and outside flakes. We refer to this data-analysis protocol for heterogeneous clusters as GSER Average. These steps yield the within-flake and outside-flake dynamic moduli as revealed by 200 nm and 1 μm diameter probes, for each of the 3 reconstituted HBE samples, in Figure 7. For comparison, Figure 8 shows results using GSER average and the two alternative applications of the GSER, always applying the methods post clustering, namely averaging within-cluster MSDs and then applying GSER, and averaging within-cluster classifiers and then applying GSER. As shown in the synthetic datasets in Figure 2, the three methods yield equivalent results for homogeneous clusters, but not for heterogeneous clusters.

## Analysis of Dynamic Moduli

The storage and loss moduli for each cluster reported in Table 1 is displayed in Figure 7, using the data-analysis protocol above. Since this involves weighted averages over power law functions *ω*^*α*^, *there is a potential for within-flake and outside-flake sol-gel frequency transitions*. Due to evidence of heterogeneity in each cluster, we also display the mean ± two standard deviations of *G*^′^ and *G*^′′^ for each cluster in which there was more than one bead.

We find that for the 1μm 60% HBE sample, one within-flake cluster reflects a sol rheology and the other a gel rheology over all frequencies. The 200 nm 60% sample has clusters at the transition between gel-like and sol-like; one “cluster” contains a single bead and thus there are no intervals around the moduli curves. These results comport with the stochastic phase separation process that produces flakes.

The 1μm 80% HBE sample shows two clusters with a sol rheology, one with a gel rheology, and once again hovering around the transition. However, unlike the previously discussed sample, the cluster at the gel-sol boundary reflects a slight sol-gel transition as frequency increases. This cluster is the only one to reflect a sol-gel transition. If one averages over the entire within-flake bead ensemble, again averaging in frequency space as the only justifiable average, Figure 8, then a sol-gel transition arises due to a spectrum of beads conveying α above and below .5.

Figure 8 shows the dynamic moduli curves from all beads in the “flake-like” clusters for the 60% and 80% reconstituted HBE bulk samples. Due to non-homogeneity of the clusters, only (unjustifiable) ensemble averaging of classifiers gives a power law with linear scaling in loglinear coordinates, while both the (justifiable) GSER Average and (unjustifiable) MSD average each display a departure from power-law scaling. *An important result of the justifiable GSER Average method is evidence of a sol-gel transition from the ensemble of probes within flakes*.

While one can focus on details revealed in Figure 8, it is difficult to make strong conclusions due to low sample sizes and the underlying stochasticity of the phase separation process of flake formation.

## Concluding Remarks

Experiments and statistical methods are developed and applied to analyze particle-tracking data from reconstituted airway mucus samples that replicate changes in the MUC5B/MUC5AC ratio characteristic of CF progression. This task of particle tracking to infer equilibrium rheological properties is challenging in highly heterogenous soft materials in general, and more so with flake-burdened CF mucus where within-flake and outside-flake domains are a result of a poorly understood, stochastic, phase-separation process. In our analysis, three levels of heterogeneity are identified and characterized from the tracked bead data: (1) a coarse scale, binary separation of dense, phase-separated, mucus flakes and dilute solution of remaining mucins; (2) within flakes, a binary separation of dense domains where bead signals can versus cannot be confidently disentangled from the experimental noise floor; and (3) in light of failed tests for sample homogeneity based on tracked beads within and outside of flakes, cluster analysis is applied to the ensembles of 200 nm and 1 μm diameter beads both within and outside flakes.

*For each tracked bead*, we develop a *two-parameter classifier metric* (*α, D*_*α*_), where *α* is the power law and *D*_*α*_ is the pre-factor of the mean-squared-displacement (MSD), *of the denoised entire experimental time series*. This fractional Brownian motion (fBm) classifier of the pure medium-induced signal for each tracked bead has been shown to be a robust statistical metric for passive particle tracking in HBE mucus (cf. [6,9,10-12,14-16]). For beads in flakes, however, previous statistical metrics for fitting fBm to increment time series data are inaccurate, illustrated herein with synthetic data of noisy bead time series representative of tracked beads in dense flakes.

The fBm classifier metric involves two steps, each utilizing the full bead time series. The first *predictor step* is based on a least-squares fit to the mean-squared-displacement of fBm, *MSD*_*X*_(*nτ*) = 2*dD*_*α*_(*nτ*)^*α*^, where *nτ* are all experimental lag times and *τ* is the minimum lag time between recorded bead positions of the microscope. This yields an initial (*α, D*_*α*_) predictor estimate, which stabilizes the *corrector step* using the fARMAs method initially developed in and extended for this study in [33]. This predictor-corrector method is shown to accurately recover the truth in synthetic data mimicking beads in flakes for which previous metrics fail.

We next apply cluster analysis to assess heterogeneity in each of the three reconstituted samples of flake-burdened mucus reflected by each bead diameter. While the power law *α* is dimensionless, the MSD pre-factor *D*_*α*_ has fractional, *α*-dependent time units. We transform *D*_*α*_ for each bead to have either the same physical units or a common nondimensionalization. We show multiple ways to do this, each giving similar clustering results using a standard clustering algorithm. We emphasize that clustering is implemented on the denoised time series classifier of the data. Once clustering is performed, we test homogeneity of the clusters to justify use of either the single bead or ensemble-averaged GSER to infer dynamic moduli of all clusters within and outside flakes for each bead diameter. Clusters within and outside flakes invariably fail a homogeneity test, strongly supporting the rheology protocol for each identified cluster: application of GSER for each single bead, and then frequency-space averaging of *G*^′^ and *G*^′′^ over all beads in the cluster. This protocol gives the dynamic moduli of inside-flake and outside-flake clusters for each of the three samples from both bead diameters. We find that flakes possess both sol and gel domains while the remaining dilute mucin solution is sol-like, both nonhomogeneous, consistent with the stochastic phase-separation process generating the flake-burdened mucus samples.

The fBm classifier (*α, D*_*α*_) has an additional advantage. For each individual tracked bead, the fBm classifier yields an exact power law for the mean-squared-displacement (MSD).

Furthermore, the Fourier transform of a power law function is also exact and a power law function, so the GSER applied to the classifier of each bead yields *an exact power-law formula for the complex modulus, G*^*∗*^(*ω*), avoiding numerical approximation. Since all within-flake and outside-flake experimental clusters fail the test for homogeneity, within-cluster averaging is performed in the frequency domain post application of the GSER to the denoised MSD for individual beads in each cluster. *This results in a non-power law rheology both for each cluster within both flakes and for the remaining dilute solvent*. The within-flake data with both 200 nm and 1 μm diameter beads in samples from the same reconstituted batch also reveals *probe size-dependent heterogeneity*. Finally, we use the relative number of 200 nm vs. 1 μm diameter beads that enter and reside in flakes to *roughly estimate flake pore size distributions to reside predominantly between 200 nm and 1* μm, *with some percentage of pores above 1* μ*m*. Further experiments and analysis are necessary to more quantitatively estimate pore-size distributions.

## Supplemental Material

### The fARMAs Denoising Method

For ease of presentation, we describe the method for a one-dimensional particle trajectory X(t), with extension to higher dimensions in [15]. The fARMA model [15] assumes that *X*(*t*) is fractional Brownian motion with linear drift,

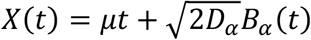

However, we observe not *X*(*t*) but rather *Y*_0_, …, *Y*_*N*_, corresponding to noisy measurements of *X*_*n*_ = *X*(*n* · *τ*), from which we must extract the pure entropic fluctuation signal. This approach to signal extraction was first proposed by Savin and Doyle [27] using the measurement model

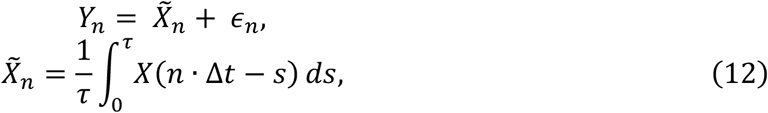

where *∈*_*n*_ is white noise representing “static” localization errors, and 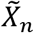 represents “dynamic” errors due to motion of the particle during the camera open aperture time *τ*. Static errors have the effect of raising the MSD of *Y* relative to that of *X* at the shortest timescales, whereas dynamic errors have the opposite effect. While theoretically appealing, Ling et al. [15] found the Savin-Doyle model lacking in flexibility to account for the complex interactions between signal and noise from one experimental lab to another. Instead, they propose an ARMA model (fARMA),

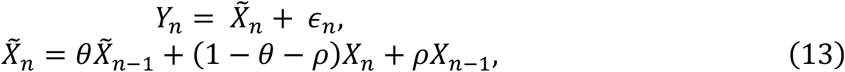

to which we have added the static noise term *∈*_*n*_ here and in [33] that we have found necessary for in-flake particles, illustrated in Figure 3 with synthetic data. We refer to model (13) as fARMAs.

### Parameter Estimation

To estimate the parameters of the fARMAs model (13), it is convenient to transform to an unconstrained basis; this improves model convergence around the parameter endpoints in the standard basis. Since *α ∈* (0, 2), an intuitive unconstrained transformation is:

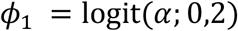

where the generalized logit transformation is given by:

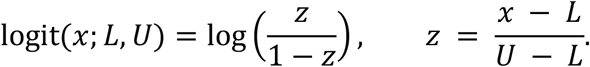

Similarly, the noise parameters are transformed using the logit transform: for the fARMAs model

*ϕ*_2_ = logit(*ρ*; −1,1) and *ϕ*_3_ = logit(*θ*; −1,1). The drift coefficients, *μ*, are inherently unrestrained, so the last parameter to transform is the scaling matrix Σ. The unconstrained parametrization for Σ is:

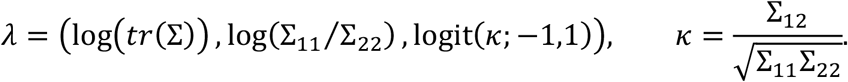

Optimization is then done using maximum likelihood estimation (MLE). However, MLE requires numerical optimization of the likelihood function, which can be highly sensitive to the choice of initial value. To address this, we develop a predictor-corrector algorithm: the predictor calculates the LS estimates of (*α, D*_*α*_), the corrector uses predictor estimates to initialize the fARMAs model.

### Choosing a threshold for the Noise Floor

Beads that become embedded in mucosal flakes exhibit low mobility. At the lowest levels of mobility, *α* < 0.1, fARMAs is unstable, especially in the estimation of *D*_*α*_ as shown with beads stuck to the glass plate in Figure 9.

## Accuracy Comparison of fARMAs and LS on Truth Set Data

The accuracies of LS and fARMAs are tested using a set of synthetic truth sets, using 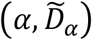 values consistent with experimental data. Two high-frequency error models are included in the simulation: no error, and static and dynamic error. In addition, linear drift is added to all simulated trajectories. Six pairs of 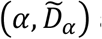 are used to simulate trajectories: one for water, two for in-between water-like and flake-like, two flake-like, and one in the noise floor. One hundred trajectories are simulated for each set of (*α, D*_*α*_) values: *α* = (0.999, 0.75, 0.5, 0.25, 0.15, 0.05),

*D*_*α*_ = (0.43, 0.108, 0.0108, 0.0027, 0.0014, 0.00096) pulled from experimental data.

For each set of values one hundred trajectories are simulated using pure fBm and then fBm + static/dynamic noise. For both methods, constant linear drift is added. For *X* and *Y* the drift coefficient is chosen using the runif() function in R. This generates a random number from a uniform distribution, in our case between -0.025 and 0.025.

Trajectories are simulated from the fBm autocorrelation function, fbm_acf() in the **subdiff** package [37], and then converted to increments using the rnormtz() function in the **SuperGauss** package [38]. For the fBm + noise model, trajectories are simulated via the same process but using fsd_acf(), the Savin and Doyle error model autocorrelation function, instead of fbm_acf(). For all (*α, D*_*α*_) pairs the value of *τ* = 0.5 is used, while the signal to noise ratio changes: *SNR* = (10, 5, 1, 0.1, 0.09, 0.08).

Once a trajectory is simulated, LS is used to get an estimate, which is used as the predictor step to be corrected by both fARMA and fARMAs models. To determine which model returns the most accurate estimate, the mean squared error, MSE, is computed between the fitted MSD and the empirical MSD between 1/60 and 10 seconds. Whichever model gives the smaller MSE is then used, and both the LS and fARMA(s) model’s *α* and *D*_*α*_ values are recorded. Figure 11 shows the true values and mean estimates (left) and then error bars corresponding to twice the mean standard error for each (right).

Figure 11 shows that in cases of no high frequency noise or when α and 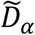 are close to water values, LS and fARMAs give similar results. However, when there is high-frequency error in the signal and α-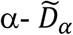 exhibit a flake-like signal, fARMAs corrects inaccuracy of LS when the LS predictor-fARMAs corrector method is used. For signals in the noise floor and with highfrequency error, both LS and LS-fARMAs are inaccurate.

## Data Cleaning

Before trajectories are analyzed, the data is cleaned. First, we filter non-isolated beads -- those within five diameters of another bead at the initial time step. This is necessary as our classifier approach assumes beads are undergoing fluctuations solely due to medium fluctuations, whereas non-isolated beads transmit forces to one another [17, 18, 19], discussed further below. The second filter ensures trajectories contain the full increment time series, automatically enforced in our experimental lab. For data in this paper, results of data cleaning are given in the table below.

| Sample | Total Beads | Proximity Filter | Length Filter | Total Left |
| --- | --- | --- | --- | --- |
| 1μm HBE + Calu3 60% | 154 | 29 | 0 | 125 |
| 1μm HBE + Calu3 80% | 137 | 2 | 0 | 135 |
| 1μm HBE + Calu3 90% | 36 | 2 | 0 | 34 |
| 200nm HBE + Calu3 60% | 17 | 0 | 0 | 17 |
| 200nm HBE + Calu3 80% | 31 | 0 | 0 | 31 |
| 200nm HBE + Calu3 90% | 32 | 0 | 0 | 32 |

| Sample | Water-Like signals | Flake-Like signals | Noise Floor signals |
| --- | --- | --- | --- |
| 1μm HBE + Calu3 60% | 89 | 18 | 18 |
| 1μm HBE + Calu3 80% | 101 | 23 | 11 |
| 1μm HBE + Calu3 90% | 23 | 8 | 2 |
| 200nm HBE + Calu3 60% | 0 | 6 | 11 |
| 200nm HBE + Calu3 80% | 3 | 21 | 1 |
| 200nm HBE + Calu3 90% | 0 | 0 | 30 |

| Sample | Percent Water-Like | Percent Flake-Like | Percent Noise Floor |
| --- | --- | --- | --- |
| 1µm HBE + Calu3 60% | 71% | 14% | 14% |
| 1µm HBE + Calu3 80% | 74% | 17% | 8% |
| 1µm HBE + Calu3 90% | 69% | 24% | 6% |
| 200nm HBE + Calu3 60% | 0% | 35% | 64% |
| 200nm HBE + Calu3 80% | 12% | 84% | 4% |
| 200nm HBE + Calu3 90% | 0% | 0% | 100% |

Beads inside flakes, especially for 1 μm beads in the 60% solution, may be within five diameters of one another, an inherent effect of flake size relative to the beads. The fit of each bead time series to pure fractional Brownian motion (fBm) plus high and low frequency noise, and insertion of the MSD formula from the inferred fBm classifier parameters into the generalized Stokes-Einstein relation, assume the only forces acting on the bead are from fluctuations of the medium. However, beads in close proximity propagate forces between one another. This coupling is the basis for two-bead microrheology (cf. [1, 3, 17, 19, 20, 18]), which we do not apply here. For all remaining bead trajectories after the first two filters, we estimate the diffusion parameters, *α* and *D*_*α*_. A final filter is then applied to bead trajectories indistinguishable from the noise floor of this experimental setup. The remaining bead data after these three filters is then analyzed.

## Acknowledgements

Partial financial support for this work is provided by the Canadian NSERC Discovery Grant RGPIN-2020-04364 for ML; NSF CISE-1931516 for MGF; Sloan Foundation G-2021-14197 for MJP, RF and MGF; the Cystic Fibrosis Foundation FREEMA19G0 for MJP and RF; the Cystic Fibrosis Foundation (HILL19G0, HILL20y2-OUT, BOUCHE19R0) and the National Institutes of Health (P30DK065988 and 1P01HL164320) for DBH and MM.

## Author Declarations

## Ethics Approval Statement

Not relevant.

## Author Contributions

**Neall Caughman:** Writing – original draft (equal); software (supporting); formal analysis (equal); writing – review and editing (equal). **Greg Forest:** Conceptualization (equal); writing – original draft (equal); formal analysis (supporting); writing – review and editing (equal). **Ronit Freeman:** Conceptualization (equal); experimental methodology (equal); writing – review and editing (equal). **David Hill:** Experimental methodology (equal); writing – review and editing (supporting); formal analysis (supporting). **Martin Lysy:** Software (lead); formal analysis (equal); writing – review and editing (equal). **Matt Markovetz:** Experimental methodology (equal); writing – review and editing (supporting); formal analysis (supporting). **Micah Papanikolas** Conceptualization (equal); experimental methodology (equal); writing – review and editing (equal).

## Data Availability

